# Resting-state EEG signatures of Alzheimer’s disease are driven by periodic but not aperiodic changes

**DOI:** 10.1101/2023.06.11.544491

**Authors:** Martina Kopčanová, Luke Tait, Thomas Donoghue, George Stothart, Laura Smith, Aimee Arely Flores Sandoval, Paula Davila-Perez, Stephanie Buss, Mouhsin M. Shafi, Alvaro Pascual-Leone, Peter J. Fried, Christopher S.Y. Benwell

## Abstract

Electroencephalography (EEG) has shown potential for identifying early-stage biomarkers of neurocognitive dysfunction associated with dementia due to Alzheimer’s disease (AD). A large body of evidence shows that, compared to healthy controls (HC), AD is associated with power increases in lower EEG frequencies (delta and theta) and decreases in higher frequencies (alpha and beta), together with slowing of the peak alpha frequency. However, the pathophysiological processes underlying these changes remain unclear. For instance, recent studies have shown that apparent shifts in EEG power from high to low frequencies can be driven either by frequency specific periodic power changes or rather by non-oscillatory (aperiodic) changes in the underlying 1/f slope of the power spectrum. Hence, to clarify the mechanism(s) underlying the EEG alterations associated with AD, it is necessary to account for both periodic and aperiodic characteristics of the EEG signal. Across two independent datasets, we examined whether resting-state EEG changes linked to AD reflect true oscillatory (periodic) changes, changes in the aperiodic (non-oscillatory) signal, or a combination of both. We found strong evidence that the alterations are purely periodic in nature, with decreases in oscillatory power at alpha and beta frequencies (AD < HC) leading to lower (alpha + beta) / (delta + theta) power ratios in AD. Aperiodic EEG features did not differ between AD and HC. By replicating the findings in two cohorts, we provide robust evidence for purely oscillatory pathophysiology in AD and against aperiodic EEG changes. We therefore clarify the alterations underlying the neural dynamics in AD and emphasise the robustness of oscillatory AD signatures, which may further be used as potential prognostic or interventional targets in future clinical investigations.

## 1. Introduction

Alzheimer’s Disease (AD) is the leading cause of dementia, accounting for 60-80% of all cases (Khan, 2016). According to the World Health Organization (2022), there are 55 million people worldwide living with dementia, while the number is estimated to reach 139 million by 2050. Consequently, it poses great economic and health challenges. Pathologically AD is primarily characterised by accumulation of amyloid-beta plaques and neurofibrillary tangles which are associated with synaptic loss and cortical volume loss. Memory impairment is one of the earliest clinical symptoms, and this progressively generalises across a wide range of cognitive domains (Jack et al., 2013). However, evidence shows that pathophysiological changes precede the appearance of clinical symptoms by decades, thus complicating attempts to avert or slow disease progression (Khan, 2016). Additionally, current early biomarkers such as the presence of amyloid-beta are not strongly predictive of cognitive function, thus complicating the estimation of cognitive decline (Blasko et al., 2008). This highlights the need for estimation of cognitively relevant neural function during the early stages of the disease that could provide prognostic markers of AD progression (Jack Jr. et al., 2018) and guide therapeutic interventions before irreversible neurophysiological and cognitive damage occurs.

A tool that has shown potential for identifying novel early-stage biomarkers of AD is electroencephalography (EEG). EEG is a non-invasive method of recording the electrophysiological dynamics of the brain, primarily generated by post-synaptic currents that are synchronous among a mass of neurons (Olejniczak, 2006). As a candidate technique for identifying biomarkers of neurological disorders, EEG has many advantages. It is non-invasive, portable, and relatively cost-effective compared to other functional neuroimaging methods such as functional magnetic resonance imaging (fMRI) and positron emission tomography (PET) (Cohen, 2017). Furthermore, neurophysiology departments trained in EEG are widely utilized in healthcare systems world-wide (Smith, 2005), meaning EEG systems and trained healthcare staff are readily available in the clinical setting.

There is growing evidence that specific EEG signatures are associated with AD, which could potentially be used to develop early-stage neurophysiological biomarkers (Horvath et al., 2018; Poil et al., 2013; Rossini et al., 2020; Tait et al., 2020). The most consistent finding comes from spectral analysis, in which the EEG signal is decomposed into its constituent frequency bands including delta (1–4 Hz), theta (4–8 Hz), alpha (8–12 Hz), beta (15–30 Hz), and gamma (30–90 Hz). Many studies have shown a ‘slowing’ of the EEG signal (Besthorn et al., 1997; Dauwels et al., 2010) characterised by power increases in lower frequencies (delta and theta) and decreases in higher frequencies (alpha and beta), along with reduction of the peak alpha frequency, in AD patients compared to healthy controls (HC) (Babiloni et al., 2004; Benwell et al., 2020; Brenner et al., 1986; Meghdadi et al., 2021; Neto et al., 2016; Schreiter-Gasser et al., 1994; Tait et al., 2019). This pattern of changes, which can be captured in a single metric by calculating the (alpha + beta)/(delta + theta) power ratio, has also been found, to a lesser degree, in conditions associated with increased risk for developing AD such as type-2 diabetes mellitus (Benwell et al., 2020; Cooray et al., 2011) and mild cognitive impairment (MCI) (Baker et al., 2008; Meghdadi et al., 2021). In AD, this power ratio has been shown to correlate with the degree of cognitive impairment (Benwell et al., 2020), thus highlighting its potential to index disease severity.

However, standard EEG spectral power analyses involve transformations that do not account for characteristics of the EEG signal that have also been found to carry functional significance. Recent methodological developments show that non-oscillatory (i.e. aperiodic) activity, which is superimposed with the periodic activity in the raw signal, can confound estimates of frequency band power (Donoghue, Haller, et al., 2020; Donoghue et al., 2021; Gerster et al., 2022; He, 2014). The neural power spectrum, which represents the amount of power across frequencies, reflects not only oscillatory activity but also the aperiodic 1/f component (Donoghue, Dominguez, et al., 2020). Oscillatory activity can be characterised by peak center frequency, the power over and above the aperiodic component, and bandwidth, whereas the aperiodic component can be measured by the aperiodic offset and exponent (Figure 1A). Changes in the aperiodic component can alter power in the band(s) of interest and be misinterpreted as oscillatory power changes (Donoghue, Dominguez, et al., 2020). Therefore, the changes in the power ratio (low frequency power increase and high frequency decrease) observed in the EEG profiles of AD patients may not reflect true changes in the periodic signal but rather changes in aperiodic features (or a combination of both). For example, a recent study of brain maturation found that controlling for the aperiodic components reversed the previous finding of decreasing alpha power from childhood to adolescence (Tröndle et al., 2022), whilst slowing of the EEG signal induced by electroconvulsive therapy has been shown to be better explained by an increase of the aperiodic exponent of the signal than by changes in oscillatory power (Smith et al., 2022). Moreover, aperiodic features themselves have been functionally linked to both healthy aging (Cesnaite et al., 2022; Merkin et al., 2021; Voytek et al., 2015) and psychopathology (Karalunas et al., 2022; Peterson et al., 2021; Robertson et al., 2019), demonstrating their functional significance and the importance of incorporating them into the analysis of AD biomarkers.

**Figure 1.**
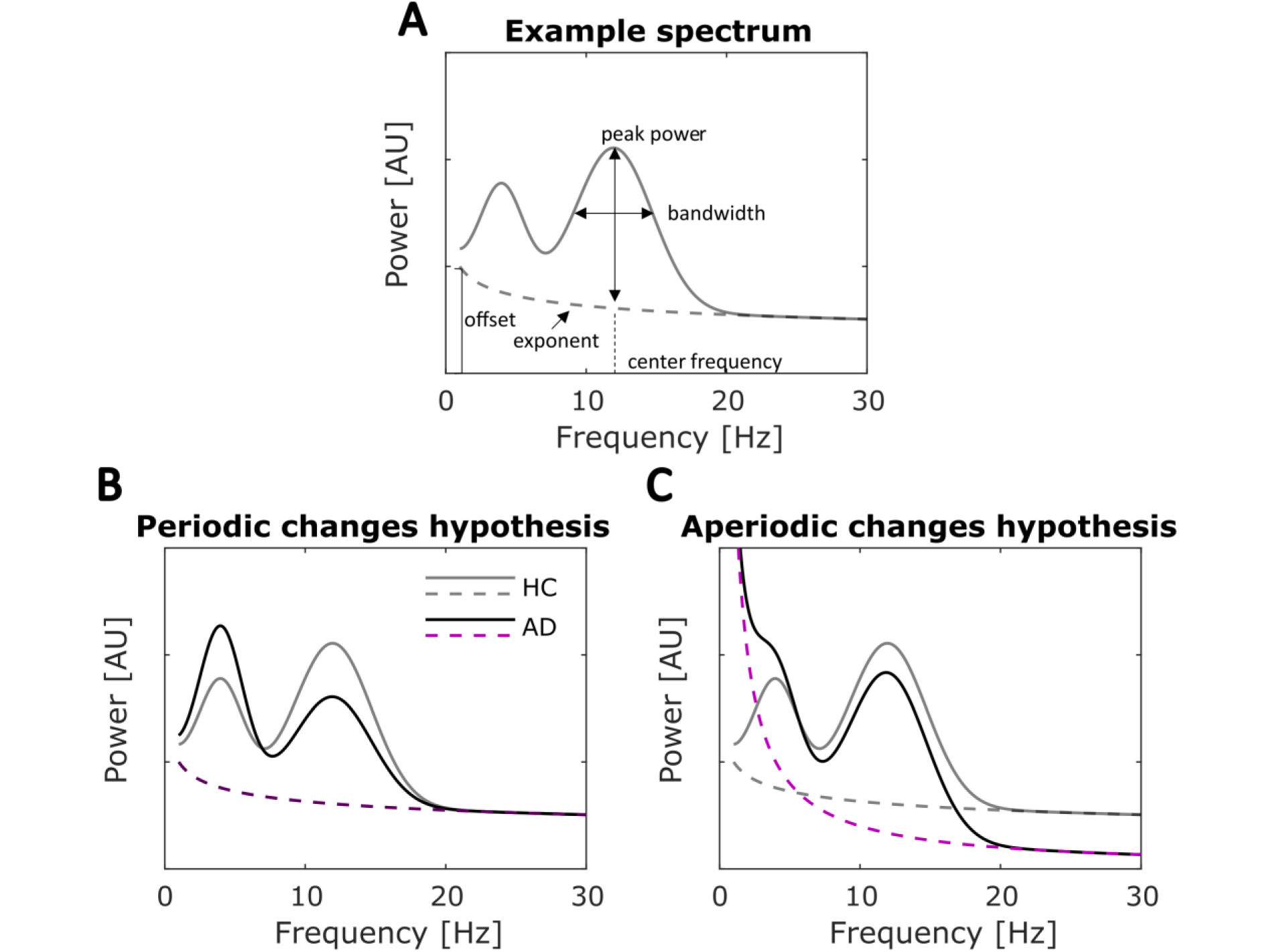
Example of a parametrised spectrum and hypothesised periodic and aperiodic changes. **A**) An example power spectrum (gray solid line) parametrised into periodic and aperiodic components (dashed line). The periodic component (over and above the aperiodic component) and each identified peak can be characterised by a peak center frequency (CF), power over and above the aperiodic component (PW), and bandwidth (BW). The aperiodic component (dashed line) is characterised by the offset (intercept) and exponent (slope) respectively. **B)** Periodic change hypothesis, with simulated data representing healthy controls (HC) in gray and AD spectra in black. The aperiodic component for HC (gray dashed line) and AD (purple dashed line) are overlapping. This panel illustrates how changes in the purely periodic activity (above the aperiodic component) can give rise to low frequency power increase and high frequency power decrease without any changes in aperiodic component. **C)** An illustration of how changes in aperiodic component (dashed lines), without any concurrent changes in the periodic activity, could result in an overall increase in spectral power at low frequencies and a decrease in higher frequencies.

Without considering the aperiodic 1/f-like component, it is impossible to elucidate the mechanism(s) underlying the most consistent EEG biomarker of AD (power ratio change). The changes may be explained by alteration(s) in the frequency and/or power of one or more neural networks subserving true periodic (oscillatory) activity (Figure 1B). Oscillatory neural activity is ubiquitous throughout the brain (Buzsáki et al., 2013), shows reliable associations with cognitive and behavioural functions (Başar et al., 2001; Thut et al., 2012; Ward, 2003), and has been proposed to play crucial functional roles including the encoding and transfer of information (Fries, 2015; Keitel et al., 2022; Singer, 2018). Alternatively, AD related ‘slowing’ of the spectra may be explained by an increase in the exponent of the 1/f-like component (see Figure 1C), which has been shown to reflect asynchronous neuronal spiking (Manning et al., 2009; Miller et al., 2014) and to index the neural excitation/inhibition ratio (Gao et al., 2017). Therefore, dissociating between these alternative mechanisms represents a crucial step towards uncovering the nature of electrophysiological abnormalities associated with AD. Here, across two independent datasets, including data previously published by Benwell et al. (2020) and Flores Sandoval et al. (2023), we separated the aperiodic EEG signal from the oscillatory signal, while also parameterizing the aperiodic features: offset and exponent (Donoghue et al., 2020) Thus, we were able to test whether EEG changes associated with AD reflect true oscillatory changes, changes in aperiodic features of the signal, or a mixture of both.

## 2. Methods

### 2.1. Study design

This study was carried out using two independent cohorts of Alzheimer’s patients and cognitively healthy controls. The data for both cohorts 1 and 2 was collected at the Berenson-Allen Center for Non-Invasive Brain Stimulation (BA-CNBS) and at Beth Israel Deaconess Medical Center (BIDMC) in Boston, MA, USA and previously published in Benwell et al. (2020) for cohort 1 and in Flores Sandoval et al. (2023) for N = 23 participants out of cohort 2.

### 2.2. Participants

Resting state EEG and neuropsychological test data were analysed from individuals who took part in research at the BA-CNBS between 2012-2020. The research was approved by local institutional ethics boards and all participants gave written informed consent prior to data collection in accordance with the Declaration of Helsinki. The following groups were included:

*Cohort 1*: A total of 45 individuals. 18 (11 females, 52-86 years old) had a probable diagnosis of mild-to-moderate AD according to DSM-5/NINCDS-ADRDA criteria (McKhann et al., 2011) with a clinical dementia rating (CDR) of 1 and a Mini-Mental State Examination (MMSE) (Folstein et al., 1975) score between 18 and 24. Six participants were on cholinesterase inhibitors, 9 on cholinesterase inhibitors and memantine, and 3 were not medicated with dementia-specific medication. Additionally, 27 healthy controls (17 female, 50-77 years old) with normal cognition (MMSE ≥ 27) and without a clinical diagnosis of diabetes (glucose metabolism HbA1c < 6.5%) were included. Additional general inclusion criteria were an age adjusted score above 80 on the 50-item Wechsler Test of Adult Reading. Note that this cohort were included in a previously published study (Benwell et al., 2020).

*Cohort 2*: A total of 44 individuals were included. 29 adults with amyloid positive early AD (13 female, 53-80 years old). Amyloid status of was determined based on [18F]-Florbetapir PET and, where PET was not available, on assessment of cerebrospinal fluid with a lumbar puncture. Further inclusion criteria for the amyloid positive early AD group were a diagnosis of mild cognitive impairment (MCI) or mild AD by a board certified neurologist according to Petersen criteria (Petersen et al., 1997) and guidelines from the National Institute of Aging and Alzheimer’s Association workgroup (N = 25), and the NINCDS-ADRDA criteria (McKhann et al., 2011) (N = 4). Additionally, 17 individuals had a CDR score of 0.5 and MMSE score ≥ 21; and 12 individuals had a CDR = 0.5-1 and MMSE ≥ 20 (Folstein et al., 1975). 15 healthy cognitively normal (MMSE ≥ 27 and CDR = 0) controls (7 female, 55-87 years old) were also included. Data from a subset of cohort 2 (N =23, 17 AD) has been published in Flores Sandoval et al. (2023).

For both cohorts 1 and 2, general inclusion criteria were an absence of other unstable medical and neuropsychiatric conditions. All participants underwent a structured neurological examination, medical history review, formal neuropsychological testing, and an EEG visit. Demographic characteristics (Supplementary Table S1), including age and education, were compared between AD and HC groups within both cohorts using independent samples t-tests, whilst non-parametric Kruskal-Wallis tests were used to compare MMSE scores. Handedness and gender proportions were also compared using Fisher’s exact test. Both AD groups were older than healthy controls, statistically significantly within cohort 1, and therefore *Age* was added as a covariate to all subsequent statistical between-group comparisons.

### 2.3. Neuropsychological testing

*Cohorts 1 and 2*: Neuropsychological testing was performed on a separate visit from the EEG recording by a trained psychometrist. Tests and inventories were drawn from the National Alzheimer’s Coordination Center’s Uniform Data Set version 1.1 (NACC-UDS) (Beekly et al., 2007) for Cohort 1, and version 3.0 for Cohort 2 (Weintraub et al., 2018). See Table 1 for tests used to examine each cognitive domain.

**Table 1.**
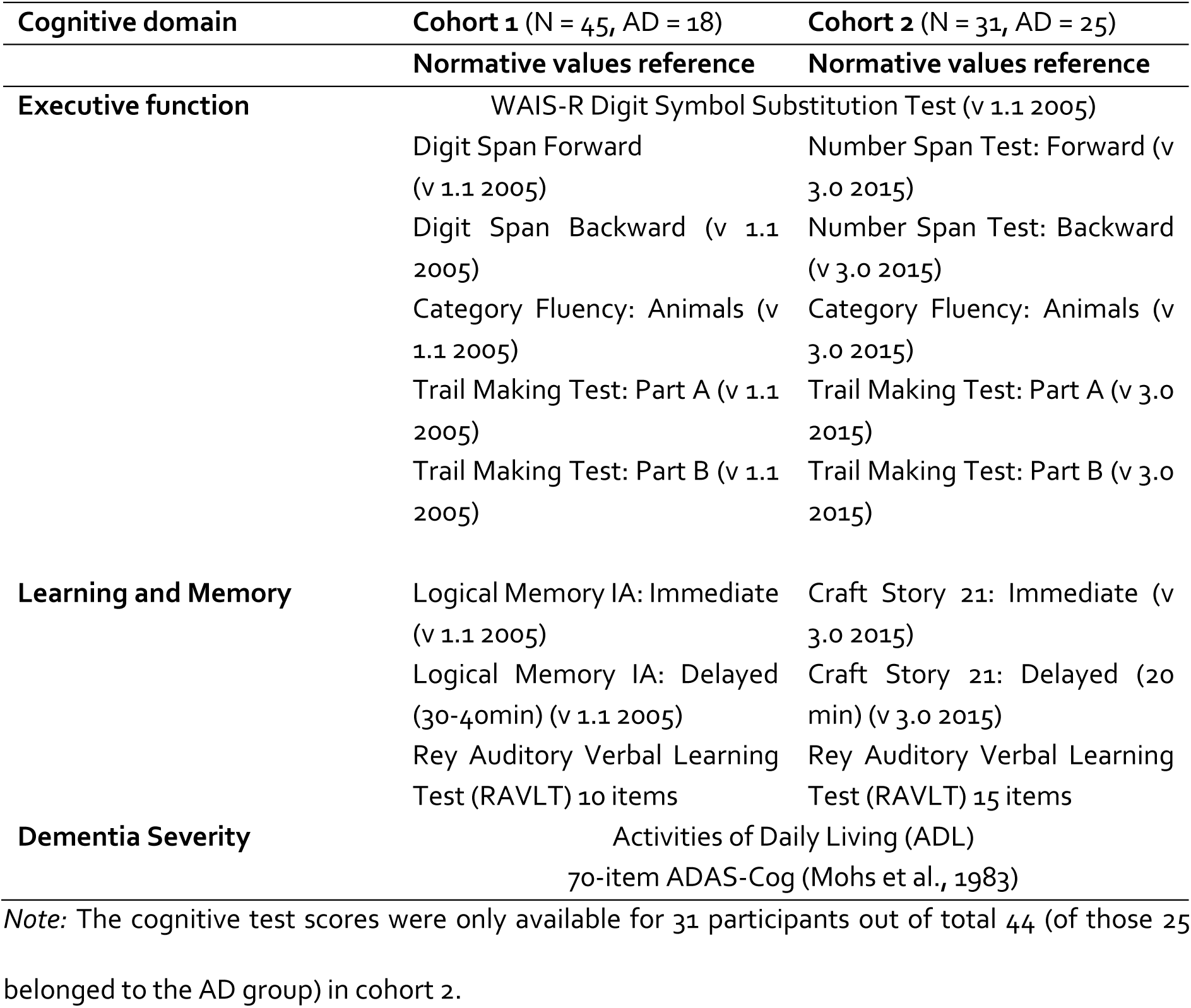
Neuropsychological tests and versions for each cohort

The raw scores for each of the above neuropsychological measures were z-scored according to normative values that were published for cognitively healthy individuals around the overall mean age across groups (Amariglio et al., 2012; Gale et al., 2007; Goldberg et al., 2010; Graham et al., 2004; Weintraub et al., 2018). Trail Making Test-A, Trail Making Test-B, ADAS-Cog Total, ADAS-Cog Recall, and ADAS-Cog Recognition scores were multiplied by -1 to ensure higher scores corresponded to better performance on all tests. Next, the z-scores from tests measuring similar cognitive functions were averaged together to form composite indices reflecting broader cognitive domains as shown in Table 1 (for similar approach see: (Benwell et al., 2020; Buss et al., 2018, 2020; Crane et al., 2012; Gibbons et al., 2012; Zadey et al., 2021). Three composite scores were computed: Dementia Severity - testing general cognitive functioning and functional independence; Executive function – testing attention, working memory, set-shifting, strategic thinking, and psychomotor processing speed; and Learning and Memory – testing verbal memory with and without context.

### 2.4. Electroencephalography acquisition and pre-processing

*Cohorts 1 and 2*: All participants underwent 5-minute resting-state EEG recordings using an extended version of the International 10-20 system with the ground and reference electrodes placed on the forehead. Two additional electrooculographic electrodes were placed at the outer canthi and below the left eye to capture horizontal and vertical eye movements. Electrode impedances were kept below 5 kΩ. The recording was obtained while participants sat in a semi-reclined chair with their eyes closed. They were instructed to remain quiet and relaxed, and to blink their eyes a few times every 1-2 minutes to maintain alertness or if observably drowsy.

The **Cohort 1** EEG data was recorded using a 64-channel system (eXimia EEG, v.3.2., Nexstim Ltd, Finland) with a sampling rate of 1450Hz. Within **Cohort 2**, data from 17 participants (10 AD) was recorded with a 60-channel system (eXimia EEG, version 3.2, Nexstim Ltd, Finland; 1450Hz sampling rate); and for 27 participants (19 AD) with a 62-channel EEG system (BrainVision, BrainProducts, GmbH, Germany; sampling rate 1000Hz). All montages were equalised by including only the 50 shared electrodes in all subsequent analyses (F1, FZ, F2, F5, F6, FP1, FP2, C1, CZ, C2, C3, C4, C5, C6,CP1,CPZ,CP2,CP3,CP4, CP5, CP6, FC1, FC2, FC3, FC4, FC5, FC6, F7, F8, FT7, FT8, TP7, TP8, TP9, TP10, T3, T4, O1, OZ, O2, P1, PZ,P2, P3, P4, P7, P8, POZ, PO3, PO4).

*Cohorts 1 and 2:* EEG data was pre-processed offline employing the same methodology and criteria for both cohorts 1 & 2 using custom written scripts in MATLAB 2016a, 2017a, and 2021a (Mathworks, USA) and incorporating EEGLAB toolbox functions (Delorme & Makeig, 2004). First, low-pass (100Hz) and high-pass (1Hz) zero-phase second order Butterworth filters were applied and a 55-65Hz notch filter was used to filter for line noise. The recordings were subsequently divided into 3-second epochs for visualisation and excessively noisy or faulty channels were removed *(M(SD)=* 2.63(2.03), range = 0-9). Next, the data were re-referenced to the average of all electrodes and noisy epochs removed using a semi-automated artifact rejection procedure *(M(SD)=* 22.09(11.60), range = 2-88). This resulted in an average of 76.5 (±9.1, range = 45-116) usable trials per participant. Independent component analysis was then run using the *fastICA* function in EEGLAB (Delorme & Makeig, 2004), and components corresponding to blinks/eye movements, muscle activity, or transient channel noise were subtracted from the data. Lastly, previously removed channels were interpolated using a spherical spline method and the data were resampled to 1024Hz.

### 2.5. Experimental design and statistical analysis

#### 2.5.1. Canonical spectral power analysis

*Cohorts 1 and 2:* The mean absolute power spectral density (PSD) across epochs was calculated for all frequency bands within the spectrum from 1 to 40 Hz at all electrodes using the *spectopo* EEGLAB function (window-size = 1024 samples, window-overlap = 512 samples, 0.1 Hz resolution) (Delorme & Makeig, 2004). The power spectra were then averaged across all electrodes for each participant and used to calculate the *original* (i.e., non-corrected) *spectra power ratio*. To enable comparisons with parametrized spectra, the lowest frequency analysed was 3 Hz (Donoghue, Haller, et al., 2020; Donoghue, Dominguez, et al., 2020; Robertson et al., 2019). Absolute power estimates within each classic frequency band (3-4Hz delta), (4-8Hz theta), (8-13Hz alpha), and (13-30Hz beta) were obtained by summing the power estimates across the frequencies contained within each band and then used to calculate an overall 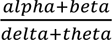 ratio for each participant.

#### 2.5.2. Spectral parameterization into periodic and aperiodic components

*Cohorts 1 and 2:* Single-participant full scalp EEG power spectra were then parameterized using the ‘Spectral Parameterization (specparam)’ toolbox (also known as ‘Fitting Oscillations One Over F’ (FOOOF) toolbox) (Donoghue, Haller, et al., 2020) in MATLAB 2021a. The ‘specparam’ algorithm separates the aperiodic EEG signal from the oscillatory signal through an iterative fitting procedure, while also parametrising the aperiodic features: offset and exponent. First, the individual PSDs were visually inspected to select the appropriate aperiodic mode reflective of the nature of the aperiodic component in log-log space (i.e., linear vs ‘knee’). Consistent with prior work (Donoghue, Haller, et al., 2020; Donoghue, Dominguez, et al., 2020; Robertson et al., 2019), a 3-40 Hz frequency range (0.1 Hz resolution) was selected for the main analyses and the spectra were fit in the ‘fixed’ (i.e., linear) mode. The remaining settings were as follows: *peak width limits* (1-12), *peak threshold* (1.0), *aperiodic mode* (fixed), *maximum number of peaks* (7), and *minimum peak height* at default (0). Additionally, the goodness-of-fit of the final model was quantified by computing frequency-wise differences between the raw spectra and the final model fits as well as computing R-squared and error metrics (Cohort 1: 𝑅^2^(error) = AD(0.997(.026)), HC(0.997(.033)); Cohort 2: AD(0.997(.029)), HC(0.996(.032)) – indicating good fits).

Both the aperiodic *offse*t and *exponent* were extracted for each participant and used in subsequent statistical analyses. The *offse*t, also known as broadband intercept, quantifies the overall up-and-down translation of the power spectrum, whilst the *exponent* is equivalent to the (negative) slope of the log-log power spectrum (in ‘fixed’ mode), with smaller *exponent* values reflecting shallower spectra. Periodic parameters of the oscillatory peak with the highest power (over and above the aperiodic component) falling within an extended alpha range (5-15 Hz) were additionally obtained, including the peak *center frequency (CF)*, *power (PW*; above the aperiodic component), and *bandwidth (BW)*.

To assess group differences in spectral power measures while controlling for aperiodic influences, aperiodic-adjusted power spectra were computed by subtracting the ‘specparam’ tool generated aperiodic fit from the raw spectra. The aperiodic-adjusted PSDs were then used to calculate the *aperiodic adjusted spectral power ratio (SPR)* 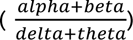, providing an estimate of periodic-only activity.

Additionally, the raw power spectra exhibited a bend in the aperiodic component (observable when plotted in log-log space) across 3-40 Hz in a subset of participants in cohorts 1 & 2. Therefore, the spectral parametrization was re-run with a model including an additional aperiodic parameter – *knee*, which captures a bend, or knee, in the shape of the power spectrum (see Supplementary Section 2 for details). This allowed us to calculate the knee frequency, which represents the estimate of the frequency at which the aperiodic component changes from horizontal to negatively sloped. We found significant between-group differences in the knee-frequency in both cohorts 1 & 2 which are reported in full in the Supplementary Material. However, while human electrophysiological recordings often show a knee (especially in higher frequency ranges) (Gao et al., 2020; Seymour et al., 2022), currently little is known about the neurophysiological significance of the knee parameter in the EEG signal, we refrain from making strong interpretations of these results (see further discussion in Supplementary Section 2).

#### 2.5.3. Statistics

Between-group differences in all EEG measures, periodic and aperiodic, were tested using separate Analyses of Covariance (ANCOVAs) with *Diagnostic Group* as independent variable, *Age* as covariate, and *EEG* as dependent variable for each of the following: *original SPR, aperiodic-adjusted SPR, offset, exponent, knee frequency* (Supplementary Section 2)*, peak power, center frequency, bandwidth*. Eta squared (𝜂^2^) was calculated as a measure of effect size. For all statistical analyses both the original and aperiodic-adjusted SPR were log transformed to normalise their distributions.

We also analysed the extent to which band ratio measures, previously reported to capture in a single variable the power shift to lower frequencies in AD (Benwell et al., 2020; Flores Sandoval et al., 2023), may be conflated with aperiodic changes and/or changes in periodic parameters beyond peak power including center frequency and bandwidth (Supplementary Section 1). Here, bivariate correlations (Pearson’s) were calculated between the *original SPR* and *exponent, offset, knee frequency* (Supplementary Section 2)*, peak power, center frequency, and bandwidth* respectively.

#### 2.5.4. Neuropsychological functions and their relationship to EEG measures

Finally, the relationships between EEG measures and cognitive functions within the cohort 1 and cohort 2 AD groups were investigated using partial correlation analyses carried out separately for the *original* and *aperiodic-adjusted ratio*, the *aperiodic exponent* (obtained from the 3-40 Hz ‘fixed’ mode analysis), and the three dominant peak parameters (Supplementary): *center frequency, bandwidth, and peak power*. Partial correlations between the outcome measures: three composite neuro-cognitive scores for *dementia severity, learning and memory,* and *executive function*, and each EEG measure were calculated while controlling for participant *Age.* All EEG measures were z-scored to facilitate comparison between different predictors.

## 3. Results

### Participant characteristics

As expected, general cognitive test scores (MMSE) were significantly lower in AD compared to HC in both cohorts. AD participants were additionally significantly older than controls in cohort 1 (t(43) = 2.642, *p* = .011), but not in cohort 2 (t(42) = 1.799, *p* = .079), hence age was included as covariate in subsequent statistical analyses. The diagnostic groups were equal in years of education, and proportions of gender and handedness in both cohorts (see Supplementary Table S1).

### The original spectral power ratio is significantly lower in AD across both cohorts

First, the full scalp averaged raw power spectra were used to calculate the 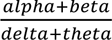 ratio for each participant. Figures 2A-B plot group averaged raw spectra for each cohort. In line with Benwell et al., (2020) and Flores-Sandoval et al., (2023), an ANCOVA testing the main effect of *Group,* while controlling for *Age,* showed the SPR was significantly higher in healthy controls compared to AD in both cohorts (Cohort 1, *F*(1,42)= 26.192, *p* < .0001, 𝜂^2^**=** .624, Figure 2C; Cohort 2, *F*(1,41)= 6.436, *p* = .0151, 𝜂^2^ **=** .157, Figure 2D). In cohort 1, this effect was driven by high frequency power changes as no significant differences were found in the low frequency power (3-8 Hz) (F(1,42) = .005, p = .942, 𝜂^2^< .001), whilst high frequency power (8-30 Hz) differed significantly (F(1,42) = 14.361, *p* < .001, 𝜂^2^ = .342 (Figure 2E&G). In cohort 2, neither low nor high frequency power alone showed significant between-group differences: 3-8 Hz: F(1,41) = .546, p = .464, 𝜂^2^ = .013; 8-30 Hz: F(1,41) = .036, p = .850, 𝜂^2^ < .001) (Figure 2F&H).

**Figure 2:**
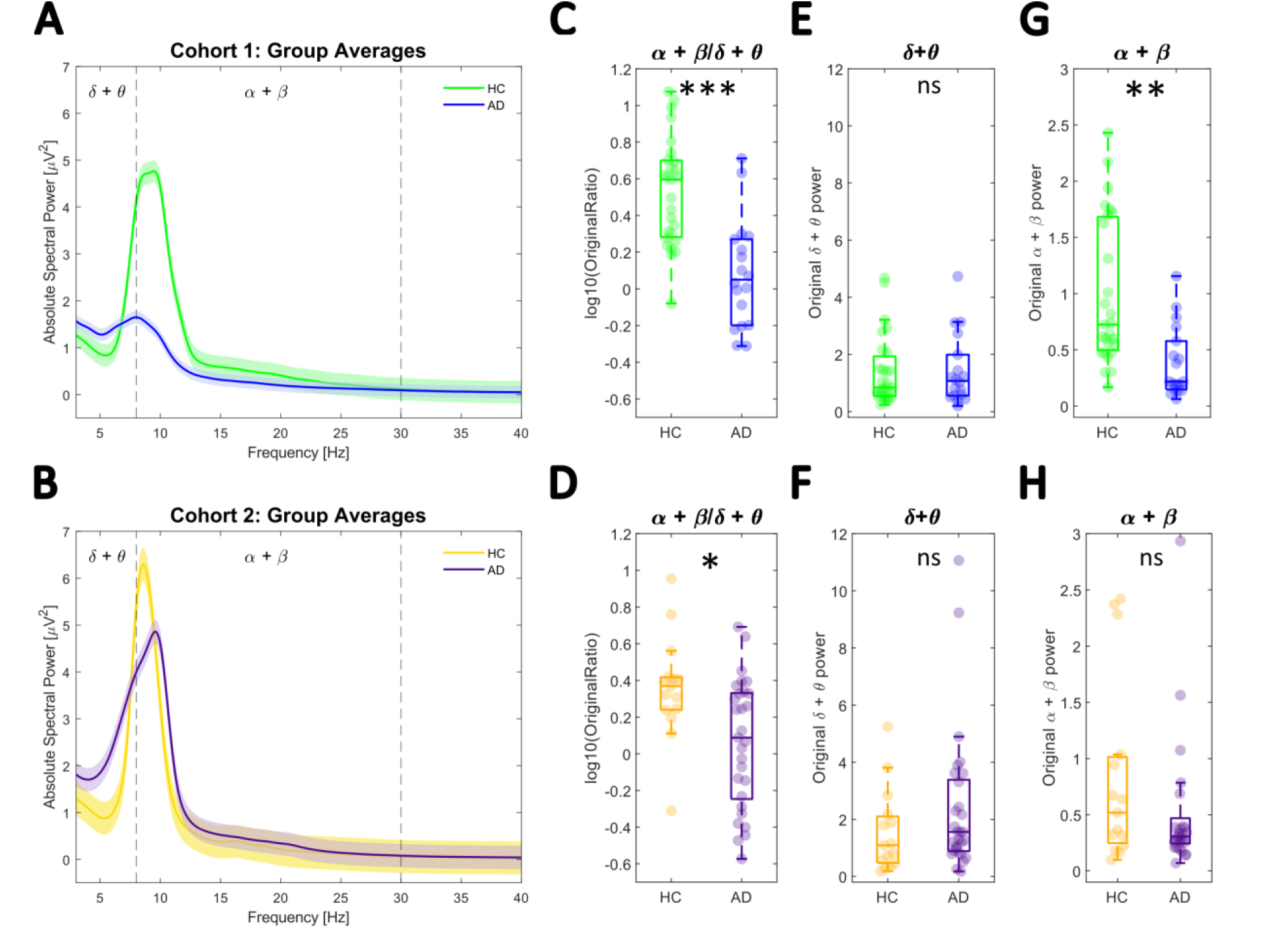
Canonical spectral power changes. **A-B** Mean full scalp power spectra for each diagnostic group. Shaded areas represent the standard error of the mean. Cohort1: green = HC, blue = AD; Cohort2: yellow = HC, purple = AD. **C-D** Comparison of the original SPR computed from the raw power spectra showed a significant difference between the groups in both cohorts. **E-F** No significant difference was found when considering low frequencies (delta + theta) alone in cohorts 1 and 2. **G-H** High frequency (alpha + beta) power increased significantly in cohort 1 but did not differ significantly in cohort 2. *** p < .0001, ** *p* < .001, * *p* < .05, ns *p* > .05.

### Spectral Parameterization

Figure 3 plots the grand average spectra for both AD and HC groups after full-scalp individual participant spectra were decomposed into periodic and aperiodic activity using the ‘specparam’ toolbox (Donoghue, Haller, et al., 2020). This allowed us to estimate both aperiodic parameters (offset and exponent) and periodic parameters (including peak power, center frequency, and bandwidth) at the individual level. Hence, we were able to investigate the respective contribution of aperiodic and periodic EEG features to the spectral power ratio and test for between-group differences. Bivariate correlation analyses confirmed that the SPR captures EEG features beyond oscillatory power alterations (Supplementary Section 1). We therefore examined AD-related changes in purely oscillatory EEG measures after controlling for aperiodic features.

**Figure 3:**
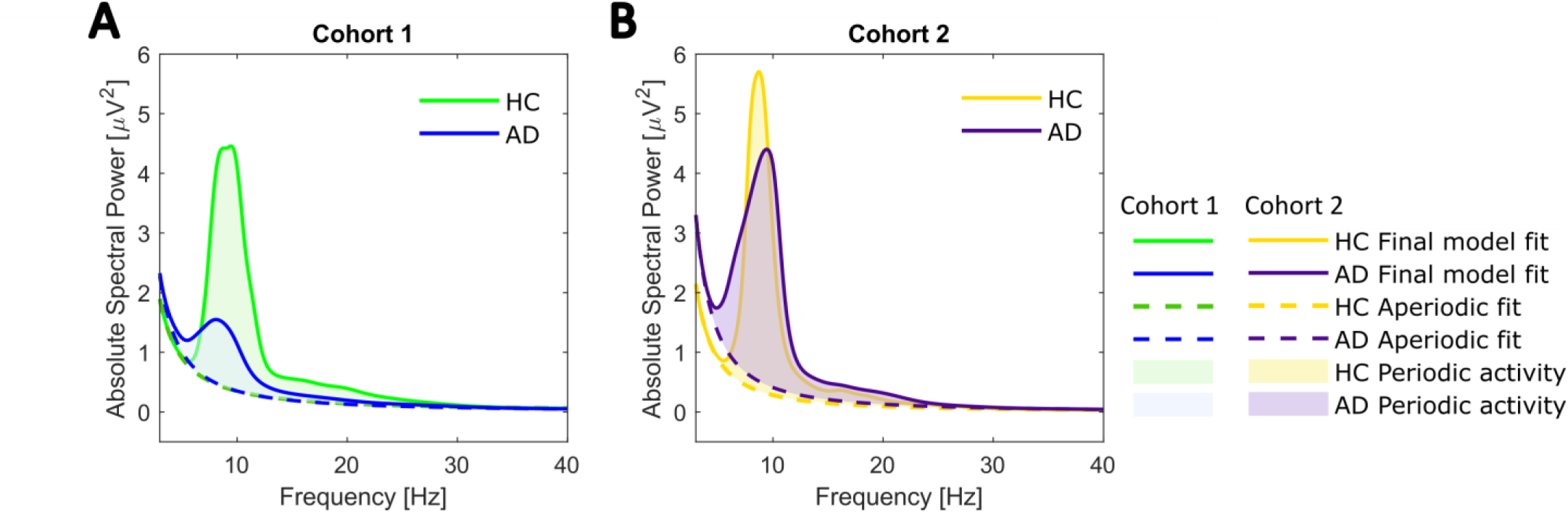
Group averages of parametrized power spectra. **A** Cohort 1. Mean full scalp power spectra for each diagnostic group after ‘specparam’ parametrization. The final ‘specparam’ model fits are in green (HC) and blue (AD). Each power spectrum further consists of periodic activity (shaded area) and the aperiodic component (dashed line). **B** Cohort 2 (yellow = HC; purple = AD).

### Periodic features (including the aperiodic-adjusted SPR) differentiate between AD and HC after controlling for aperiodic features

Figures 4A and 4C illustrate the group averaged full-scalp power spectra after removing the aperiodic component, leaving only oscillatory activity, for each cohort. We compared ‘specparam’ identified periodic parameters which characterised the dominant peak within the 5-15Hz range. Figure 4B shows that, in Cohort 1, the power over and above the aperiodic component at this peak was higher in HC compared to AD (*F*(1,42) = 24.212, *p* < .0001, 𝜂^2^ = .576) whilst the centre frequency (*F*(1,42) = 1.939, *p* = .171, 𝜂^2^ = .046) and bandwidth (*F*(1.42) = 2.862, *p* = .098, 𝜂^2^= .068) did not differ significantly. The same pattern of results was observed for Cohort 2 (Figure 4D), with peak power being significantly higher in HC compared to early AD (Cohort 2: *F*(1,41) = 4.422, *p* = .042, 𝜂^2^ = .108) and no significant differences were observed for center frequency (Cohort 2: *F(*1,41) = .597, *p* = .444, 𝜂^2^= .015) or bandwidth (Cohort 2: *F*(1,41) = 2.824, *p* = .100, 𝜂^2^= .069).

**Figure 4:**
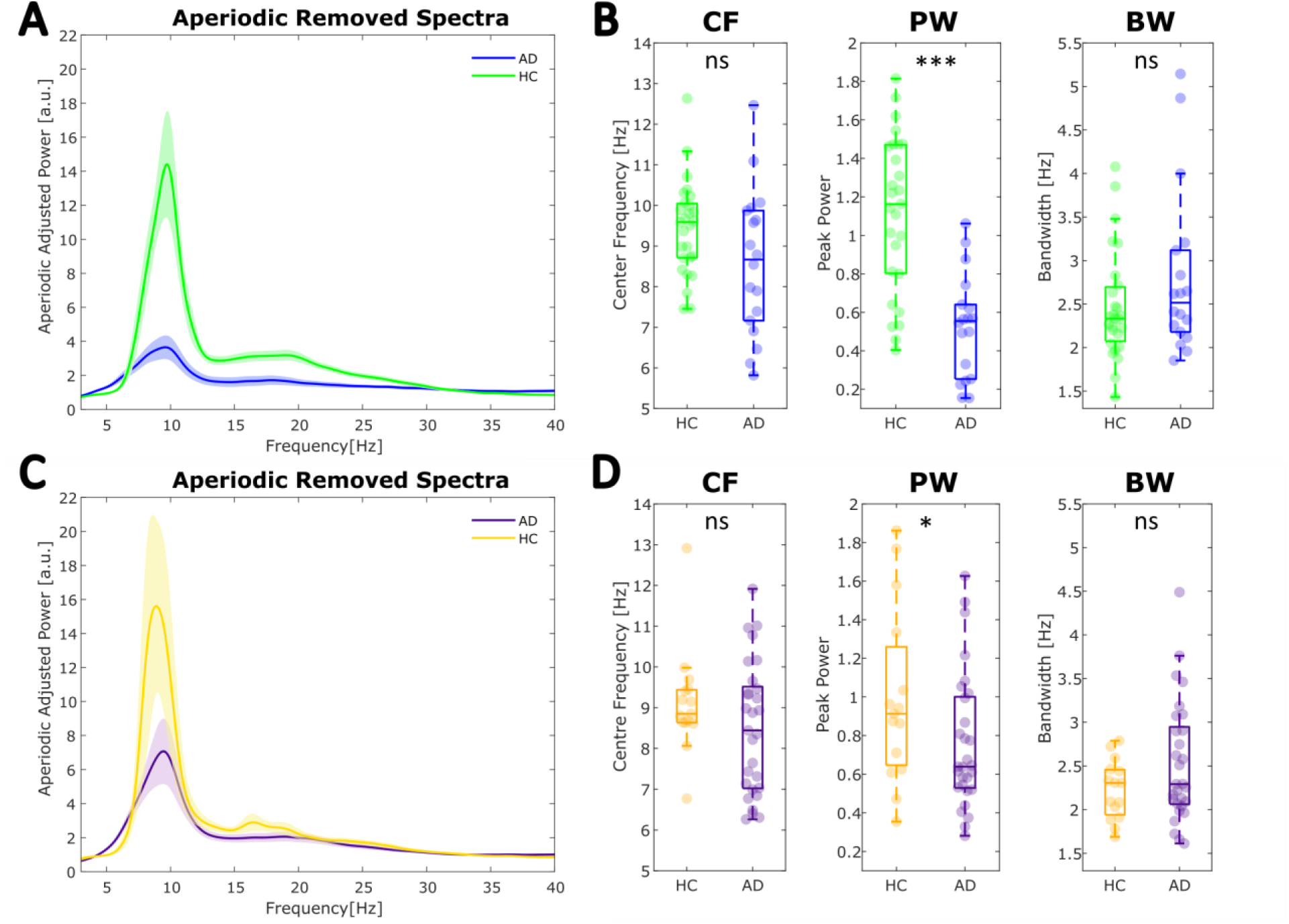
Periodic activity in full-scalp power spectra is altered in AD, exhibiting a shift in activity from higher to lower frequencies. **A-B** Cohort 1 results. **A** The group averaged spectra after the aperiodic activity has been subtracted from the raw spectra for each participant (AD: blue, HC: green). The shaded areas represent standard error. **B** Between-group comparison of periodic parameters showed power at the dominant alpha (5-15Hz) peak is significantly reduced in AD, whilst its frequency and bandwidth are not. **C-D** Cohort 2 results. **C** Group averaged periodic components of the power spectrum (AD: purple, HC: yellow). The shaded area represents standard error. **D** Peak alpha (5-15Hz) power differed between groups, whilst center frequency and bandwidth did not. CF: peak center frequency, PW: power over and above the aperiodic component, and BW: bandwidth

The aperiodic-adjusted spectra (at the individual level) were also used to compute an aperiodic-adjusted SPR (log-transformed). In line with the results of the canonical analyses (Figure 2), aperiodic-adjusted SPR was significantly lower in AD relative to HC in both cohorts (Cohort 1: Figure 5A, *F*(1.42) = 18.366, *p* < .001, 𝜂^2^= .437; Cohort 2: Figure 5B, *F*(1,41) = 5.364, *p* =.026, 𝜂^2^ = .131). Similarly to the canonical analysis results, in cohorts 1 and 2 (Figure 5C-D, E-F), the aperiodic-adjusted low frequency power (3-8 Hz) did not differ significantly between groups, whilst high frequency (8-3 0Hz) power decreased significantly in cohort 1 and showed a trend towards significance in cohort 2 (Cohort 1: 3-8Hz: F(1,42) = 0.257, p = .615, 𝜂^2^= .006 ; 8-30Hz: F(1,42) = 27.638, p < .0001, 𝜂^2^= .658; Cohort 2: 3-8Hz: F(1,41) = 0.743, p = .394, 𝜂^2^= .018; 8-30Hz: F(1,41) = 3.970, p = .053, 𝜂^2^= .097). This suggests that the oscillatory alterations in AD were primarily driven by alpha and beta power changes.

**Figure 5:**
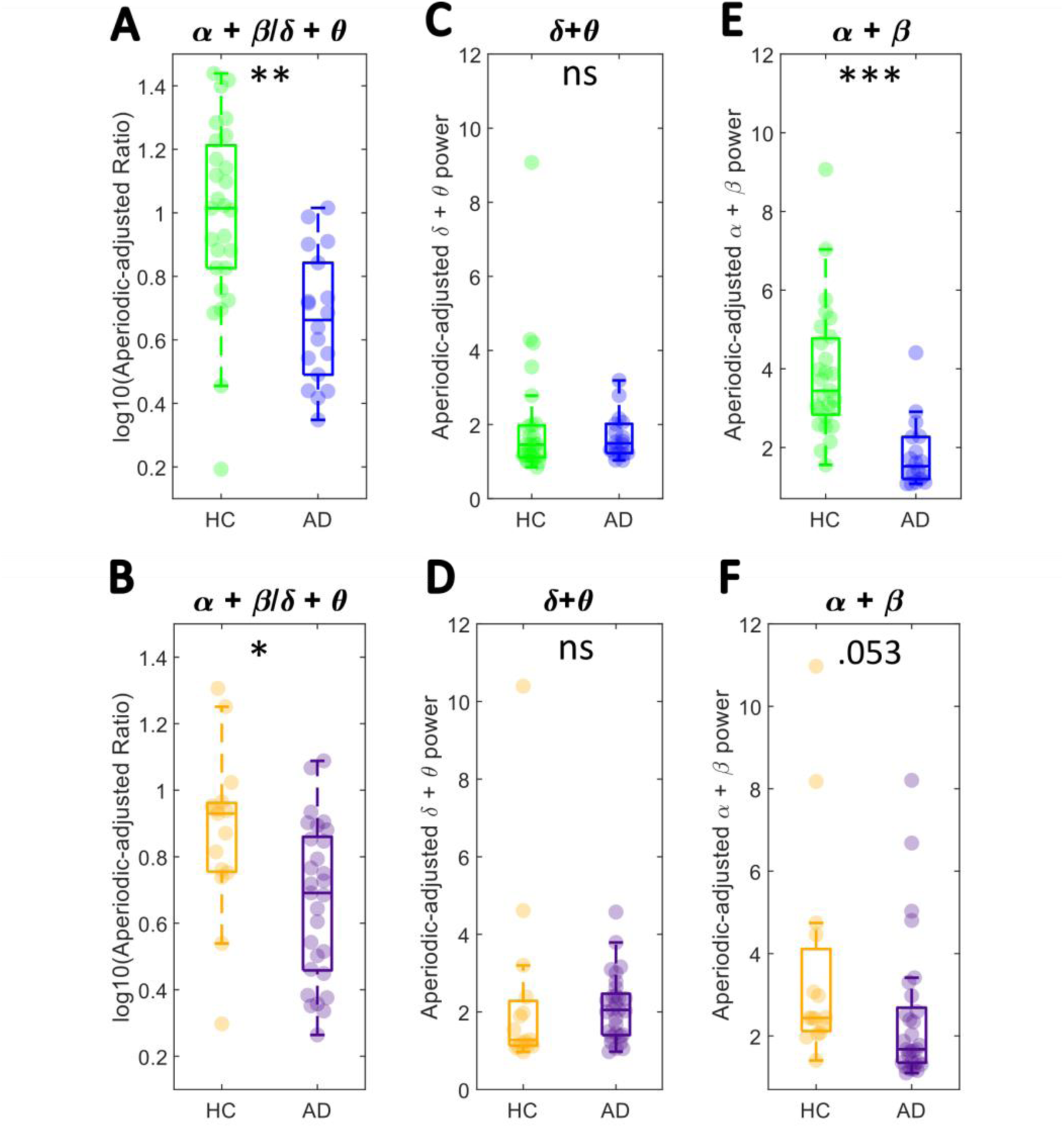
**A-B** Comparison of the aperiodic-adjusted SPR (log-transformed) showed a significant between-group difference in both cohorts. **C-D** Low frequency power (3-8 Hz) did not differ in cohorts 1&2, whilst **E-F** high frequency power (alpha + beta) decreased in AD relative to HC (this difference was statistically significant in cohort 1). *** *p* < .0001, ** *p* < .001, * *p* < .05, ns *p* > .05.

### No significant differences in aperiodic features between diagnostic groups

Figure 6 shows the results of analyses comparing the aperiodic features between AD and HC, also controlling for age, after parametrization of full scalp individual power spectra. The aperiodic 1/f components were overlapping in both AD and HC groups in Cohort 1 (Figure 6A-C) and no significant main effect of diagnostic group was found for either the offset or exponent: offset (*F*(1.42) = .114, *p* = .737, 𝜂^2^ = .0027); exponent (*F*(1,42) = .771, *p* = .385, 𝜂^2^= .018). This pattern was replicated in Cohort 2 (Figure 6D-F), with no significant between-group differences in either offset (Cohort 2: *F*(1,41) = 1.079, *p* = .305, 𝜂^2^ = .026), or exponent (Cohort 2: *F*(1,41) = .451, *p* = .506, 𝜂^2^ = .011) being found. These results, replicated across distinct cohorts, suggest AD is not associated with alteration in aperiodic EEG activity.

**Figure 6:**
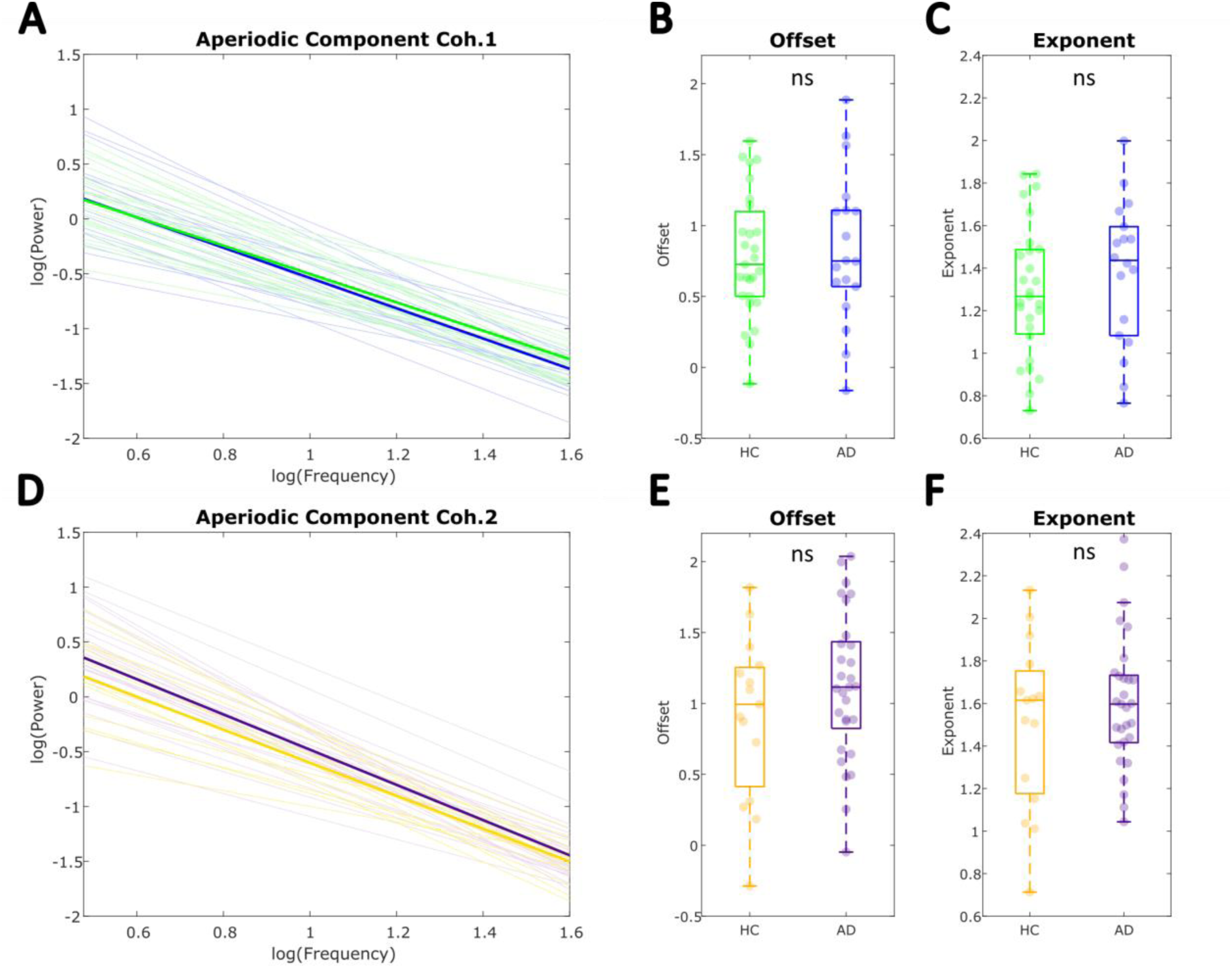
Aperiodic parameters of individual global power spectra do not differ between diagnostic groups. **A-C** Cohort 1. **A** Aperiodic component of the power spectrum (bold) averaged across individuals within each diagnostic group from Cohort 1. Individual full scalp aperiodic components are also plotted (AD = blue; HC = green). **B-C** Comparison of the aperiodic offset and exponent between AD and HC within Cohort 1 showed no significant differences after controlling for participant age with an ANCOVA (*p* > .05). **D-F** Cohort 2. **D** plots group averaged aperiodic component over individual components from each diagnostic group within Cohort 2 (AD = purple; HC = yellow). **E-F** Between-group comparisons controlling for age also showed no significant differences between HC and AD groups within Cohort 2 (*p* > .05).

### Relationships with neuropsychological function

In a final analysis, we sought to investigate the potential cognitive relevance of changes in the EEG measures of interest (original and aperiodic-adjusted SPR, aperiodic exponent and offset) in AD.

Table 2 shows the results of partial correlation analyses testing the EEG-cognition relationships while *Age* was controlled for. The original SPR significantly predicted dementia severity in both cohorts (cohort 1: p = .015; cohort 2: p = .011). When only the periodic activity was included in the SPR (aperiodic-adjusted SPR), higher SPR was significantly associated with higher dementia severity scores (p = .017) in cohort 2, while in cohort 1 this relationship did not reach statistical significance (p = .130). Higher original and aperiodic-adjusted SPR also predicted better executive function performance in Cohort 1 and 2. No other relationships were statistically significant. Importantly, these results highlight that while periodic EEG activity changes associated with AD predict disease severity, we did not find evidence of an association between aperiodic features and neurocognitive functioning in AD.

**Table 2.** Results of partial correlations (Pearson’s) for the unique relationships between periodic and aperiodic parameters and neurocognitive functions while controlling for age

| Cognitive Function | AD Group | Original Ratio | Aperiodic-adjusted Ratio | Exponent | Offset |
| --- | --- | --- | --- | --- | --- |
| | | $r(p)$ | $r(p)$ | $r(p)$ | $r(p)$ |
| **Dementia Severity** | *Cohort 1* | **0.578 (.015)** | 0.382 (.130) | -0.412 (.101) | -0.263 (.307) |
|  | *Cohort 2* | **0.512 (.011)** | **0.483 (.017)** | -0.035 (.872) | 0.168 (.433) |
| **Memory & Learning** | *Cohort 1* | 0.228 (.379) | 0.241 (.352) | -0.078 (.767) | 0.070 (.790) |
|  | *Cohort 2* | 0.255 (.228) | 0.166 (.438) | -0.018 (.935) | 0.143 (.505) |
| **Executive Function** | *Cohort 1* | **0.537 (.026)** | **0.586 (.013)** | -0.329 (.198) | -0.166 (.524) |
|  | *Cohort 2* | **0.429 (.036)** | **0.441 (.031)** | -0.100 (.641) | 0.111 (.607) |

*Note.* Each predictor was z-scored to facilitate comparability of the correlation coefficients.

These analyses were additionally run for all three periodic parameters (peak power, center frequency, and bandwidth), however, no replicable significant relationships were found between these measures and the three cognitive composites (Supplementary Table S2). Similarly, no EEG measures significantly predicted cognitive function within the healthy control group (Supplementary Table S3).

## 4. Discussion

This study aimed to clarify the physiological processes underlying the hallmark resting-state EEG marker of Alzheimer’s disease, spectral power shift from high to low frequencies, by accounting for previously overlooked EEG components. We first successfully replicated the results of the canonical spectral analyses across two independent cohorts (Benwell et al., 2020; Flores Sandoval et al., 2023). Crucially, after parametrising the EEG power spectra into periodic (oscillatory) and aperiodic activity, we found that the aperiodic-adjusted Spectral Power Ratio (SPR), which isolates changes in oscillatory power, differed significantly between AD and HC groups in both cohorts. Specifically, in comparison to HC, the AD group showed an oscillatory power decrease in high frequencies (8-30 Hz) in both cohorts, while low frequency power (3-8 Hz) did not differ significantly. In contrast, no significant differences in the aperiodic component (and its parameters: offset and exponent) were found between AD and HC. Hence, the results highlight that AD-related EEG alterations, captured in the Spectral Power Ratio, are primarily driven by periodic activity, whilst we found no evidence for aperiodic activity changes in AD. By replicating the main findings across two cohorts, we highlight the robustness of these oscillatory signatures of AD. Given the rising concerns regarding the replicability of scientific findings (Open Science Collaboration, 2015), especially within neuroscience and related fields (Button et al., 2013; Pavlov et al., 2021; Poldrack et al., 2017), the reproducibility of the findings is important.

Our results highlight that previously reported EEG signatures of AD are driven by periodic EEG activity changes, adding methodological and mechanistic clarity to previous studies. Recent work emphasises that periodic and aperiodic EEG features can be confounded when predefined frequency bands and their ratios are used (Donoghue, Dominguez, et al., 2020; Donoghue et al., 2021). Indeed, we found that the non-corrected SPR correlated significantly with aperiodic offset and exponent in the present data (in both cohorts 1 & 2: see Supplementary Section 1). Nevertheless, the SPR calculated from purely oscillatory activity differed between diagnostic groups, in line with previous findings (Benwell et al., 2020; Babiloni et al., 2004, 2016; Brenner et al., 1986; Dauwels et al., 2010; Jeong, 2004; Meghdadi et al., 2021; Neto et al., 2016; Rossini et al., 2020; Schreiter-Gasser et al., 1994; Tait et al., 2019), whereas the aperiodic features did not. These results thus lend support to an interpretation linking AD to abnormal neural oscillations relative to cognitively healthy controls. Intriguingly, this contrasts a growing body of literature showing changes in the spectral aperiodic exponent are associated with a range of neuropsychological pathologies (Pani et al., 2022), including ADHD (Karalunas et al., 2022; Robertson et al., 2019), schizophrenia (Molina et al., 2020; Peterson et al., 2021), stroke (Johnston et al., 2023), and Parkinson’s disease (Belova et al., 2021).

Additionally, the present results suggest that oscillatory abnormalities captured in the SPR are primarily driven by high frequency (8-30 Hz) power decreases. Relative to HC, we found reduced alpha + beta power (in cohort 1), as well as lower alpha peak power in AD (in cohorts 1 & 2), in line with previous literature (Babiloni et al., 2004, 2016; Benwell et al., 2020; Huang et al., 2000; Meghdadi et al., 2021). Conversely, we did not find a consistent significant slowing of the peak alpha frequency across cohorts, after controlling for the aperiodic signal, contrasting prior studies (Benwell et al., 2020; Moretti et al., 2004; Poza et al., 2007). This suggests, oscillatory power, rather than frequency, constitutes a more reliable marker of AD. Somewhat inconsistent with existing findings are our results regarding low frequency (delta + theta) activity (Babiloni et al., 2004; Benwell et al., 2020; Meghdadi et al., 2021; Moretti et al., 2004), whereby we did not observe low frequency power changes in cohort 1 or 2. This discrepancy may stem from the use of relative spectral power in previous studies that found low frequency power increases in AD (Babiloni et al., 2004; Benwell et al., 2020; Moretti et al., 2004; Tait et al., 2019). Other studies that used both relative and absolute power, only report results of statistical comparisons for relative power (Meghdadi et al., 2021). When relative as opposed to absolute power is computed, the low frequency band power is normalised by dividing it by the total power of the spectrum. Consequently, the relative power at low frequencies may also reflect any changes in power at high frequencies, potentially conflating the observed between-group differences.

Notably, the periodic nature of neural abnormalities linked to AD we observed also highlights a dissociation between healthy aging and Alzheimer’s pathology. While previous research found peak alpha frequency and power reductions in older age (Babiloni, Binetti, Cassarino, et al., 2006) that appeared to be similar to the oscillatory ‘slowing’ observed in AD, recent studies have shown that healthy aging is accompanied by a reduction in the aperiodic exponent that may have previously been confounded with oscillatory changes (Cesnaite et al., 2022; Donoghue, Haller, et al., 2020; Merkin et al., 2021; Voytek et al., 2015). For example, after controlling for age-related flattening of the aperiodic exponent, Merkin et al. (2021) found that peak alpha frequency was significantly slower in older compared to younger adults, but alpha peak power did not differ. In contrast, our results showed that AD pathology was more reliably accompanied by reductions in aperiodic adjusted alpha (+beta) power, with no alpha peak frequency slowing observed, while the aperiodic signal did not differ relative to HC in either of the cohorts. Hence, we establish that AD differentiates from the often-found aging effect on aperiodic parameters, with a periodic-specific effect found to differentiate between clinical and non-clinical participants. Interestingly, conceptually similar work examining spectral slowing in the context of stroke found influences of both periodic and aperiodic parameters (Johnston et al., 2023). These results highlight that differences in power ratios reflecting ‘spectral slowing’ can be driven by different underlying changes, and that spectral parameterizing can adjudicate between different changes. In doing so, we start to demonstrate that what might otherwise be considered similar patterns of “spectral slowing” can be shown to occur in distinct ways when differentiating between healthy aging and clinical disorders, and when differentiating between distinct disorders such as between AD and stroke.

On a physiological level, this dissociation implies that the neural alterations accompanying healthy aging may be more closely linked to abnormalities in asynchronous neuronal spiking (Manning et al., 2009) or synaptic excitation/inhibition balance (Gao et al., 2017), whilst pathological aging due to AD is more associated with aberrant neural synchrony (Babiloni et al., 2016). In particular, oscillations are proposed to reflect the global synchronisation of pyramidal cortical neurons that facilitate the integration of information from the cortex, thalamus, and the brainstem (Babiloni et al., 2016). Consequently, oscillatory abnormalities may reflect the effect of AD related neurodegeneration on large-scale cortical networks necessary for neuronal communication and cognition (Rossini et al., 2007; Uhlhaas & Singer, 2006). Interestingly, decreases in alpha power (and increases in delta + theta power) have been linked to a range of neurodegenerative processes including atrophy of the hippocampus (Babiloni et al., 2009; Fernández et al., 2003; Helkala et al., 1996), cortical gray matter (Babiloni et al., 2013), and subcortical white matter (Babiloni, Frisoni, et al., 2006). Recent studies have additionally found correlations between oscillatory ‘slowing’ and proteinopathy, such as tau decomposition (Coomans et al., 2021) and regional accumulation of amyloid-β plaques, while the strength of the latter relationship also predicted cognitive impairments (Wiesman et al., 2022).

The oscillatory nature of electrophysiological abnormalities linked to AD may additionally inform the understanding of the functional consequences of AD pathophysiology. For instance, alpha oscillations have been proposed to inversely index the degree of global cortical excitability (Romei et al., 2008). They are therefore thought to reflect functional inhibition which allows adaptive regulation of neuronal excitability necessary for facilitating top-down control and goal-directed cognitive and/or behavioural function (Jensen & Mazaheri, 2010; Klimesch et al., 2007). AD related decreases in alpha power may thus reflect a shift towards functional disinhibition relative to HC. Consequently, this may lead to cognitive impairments in domains in which top-down control plays a central role, such as executive functions. Our results are consistent with this, as a significant relationship between the SPR and executive function impairments in AD was found for both Cohorts 1 and 2, with higher SPR predicting better executive function scores.

We further found that the SPR was moderately associated with global dementia severity, even after the aperiodic signal was accounted for. This aligns with previous research linking increases in delta + theta, and decreases in alpha + beta power, to greater global cognitive impairment in AD and MCI (Babiloni, Binetti, Cassetta, et al., 2006; Claus et al., 2000; Luckhaus et al., 2008). In contrast to periodic measures, we did not find any significant relationships between aperiodic parameters and any of the neurocognitive composite scores. Hence, these results further highlight the dissociation between healthy aging and pathological changes due to AD, with the former being more tightly linked to aperiodic activity, whilst the latter is tracked by periodic EEG activity changes.

Overall, we show that AD-related changes in SPR are best explained by changes in spectral power and not by broadband changes in the EEG signal. Furthermore, the SPR has the potential to index disease severity. Notably, our results emphasize that alpha and beta periodic activity may be a particularly specific marker of AD related changes. Periodic EEG activity therefore constitutes a robust and generalisable marker of AD pathology that could be further tested for potential diagnostic and interventional applications. For instance, both invasive and non-invasive neurostimulation techniques (e.g., deep brain stimulation and transcranial direct current stimulation) have shown promising effects on the treatment of symptoms of neurodegenerative diseases including Parkinson’s and Alzheimer’s disease (Lefaucheur et al., 2017; Little et al., 2013; Scharre et al., 2018). More recently, closed-loop neurostimulation approaches, where electrical stimulation is delivered based on an ongoing electrophysiological biomarker of the pathology, have begun to be developed (Iturrate et al., 2018). The pathological changes in oscillatory activity in AD could further be investigated as candidate markers for these interventions, given their reliability and cognitive relevance. Future prospective longitudinal studies should examine whether the oscillatory abnormalities associated with AD can be used to detect individuals at risk of developing AD, as well as test their specificity to different types of dementia.

Other aspects of the present research could also be developed upon in the future. While we show robust and replicable group differences in the SPR and oscillatory power measures, additional testing of their ability to dissociate between AD and HC at an individual level is necessary to establish the suitability of oscillatory markers as prognostic tools or therapeutic targets. Additionally, we cannot completely rule out the presence of preclinical AD in our healthy control individuals. While no HCs obtained MMSE scores indicative of clinical impairment, biomarker quantification (e.g., amyloid) would be necessary to confirm the absence of AD. In addition, future studies may inspect the association between the periodic and aperiodic components of EEG activity and amyloid load.

## Conclusion

The present results advance our understanding of EEG markers of neuro-cognitive dysfunction in AD. Adding methodological and mechanistic clarity to previous work, we show that the electrophysiological spectral slowing in AD is driven by periodic oscillations, whilst aperiodic EEG features remain unaffected. Our results suggest that relative to HC, AD is most consistently characterised by high frequency (alpha + beta) power decreases and a lower SPR, with the latter also showing the potential to track disease severity. Replicated across two independent datasets, these results help uncover mechanisms underpinning changes to neural dynamics in AD. Our results lay the foundation for further investigations aimed at establishing potential diagnostic and interventional clinical applications of oscillatory AD signatures.

## Supporting information

Supplementary Section

## Acknowledgements

M.K. was supported by a Vacation Scholarship from The Carnegie Trust for The Universities of Scotland (VAC010474). C.S.Y.B was supported by the British Academy/Leverhulme Trust and the United Kingdom Department for Business, Energy and Industrial Strategy (SRG19/191169). S.S.B. was supported by the NIH (1K23AG068384-01A1) and the Alzheimer’s Association (2019-AACSF- 643094). M.M.S. was supported by the NIH (NIA P01AG031720-8405). A.P.L. is partly supported by grants from the NIH (R01AG076708, R03AG072233) and BrightFocus Foundation. The data was collected with support from the following grants: NIH-NINDS R21NS082870, NIH-NIMH R01MH115949-S1, NIH-NIA R21AG051846, and NIA R01AG060987.

## Competing interests

Dr. A. Pascual-Leone is partly supported by grants from the National Institutes of Health (R01AG076708, R03AG072233) and BrightFocus Foundation. Dr. A. Pascual-Leone serves as a paid member of the scientific advisory boards for Neuroelectrics, Magstim Inc., TetraNeuron, Skin2Neuron, MedRhythms, and Hearts Radiant. He is co-founder of TI solutions and co-founder and chief medical officer of Linus Health. None of these companies have any interest in or have contributed to the present work. Dr. A Pascual-Leone is listed as an inventor on several issued and pending patents on the real-time integration of transcranial magnetic stimulation with electroencephalography and magnetic resonance imaging, and applications of noninvasive brain stimulation in various neurological disorders; as well as digital biomarkers of cognition and digital assessments for early diagnosis of dementia.

The remaining authors declare no competing financial interests.

## Corresponding author

Correspondence to Martina Kopcanova

