## Supplementary Section for "Resting-state EEG signatures of Alzheimer’s disease are driven by periodic but not aperiodic changes"

**Supplementary Table S1.** *Participant demographics*

| Cohort | AD | HC | Statistical Comparison Cohort 1 | | |
|---|---|---|---|---|---|
| | *n* = 18 | *n* = 27 | ***df*** | ***t/χ²*** | ***p* value** |
| Age | 69.3 (7.8) | 62.4 (9.1) | **43** | **2.642** | **.011** |
| Education | 16.9 (3.9) | 15.9 (2.4) | 43 | 1.119 | .270 |
| MMSE | 21.6 (2.2) | 29.5 (0.8) | **1** | **33.599** | **<.001** |
| Female | 11 (61%) | 17 (63%) | 1 | Fisher's Exact Test, *n* = 45, 2-tailed, *p* = 1 | |
| Handedness (right) | 15 (83%) | 27 (100%) | 1 | Fisher's Exact Test, *n* = 45, 2-tailed, *p* = .058 | |
| | **AD2** | **HC2** | **Statistical Comparison Cohort 2** | | |
| | *n* = 29 | *n* = 15 | ***df*** | ***t/χ²*** | ***p* value** |
| Age | 70.6 (8.0) | 65.8 (9.1) | 42 | 1.799 | .079 |
| Education | 17.3(2.6) | 16.6 (3.1) | 35 | 0.726 | .473 |
| MMSE | 25.69(2.8) | 29.27 (0.8) | **1** | **17.443** | **< .001** |
| Female | 13 (44%) | 7 (46%) | 1 | Fisher's Exact Test, *n* = 44, 2-tailed, *p* = 1 | |
| Handedness (right) | 25 (86%) | 14 (93 %) | 1 | Fisher's Exact Test, n = 44, 2-tailed, *p* = .647 | |

*Note:* The table shows mean and standard deviation (M(SD)) for age, years of education, and MMSE scores for each cohort. Number of females and right-handed individuals is also reported. Age and years of education were compared using independent-samples t-test. Note that years of education in cohort two were only available for N = 37 (27 AD) individuals. MMSE scores were compared with a Kruskal-Wallis test, and gender and handedness were compared with Fisher's exact test.

**Supplementary Section 1: The SPR captures EEG features beyond oscillatory power alterations**

We examined the extent to which the exponent and offset of the aperiodic signal contribute to the original SPR with bivariate correlation analyses. Figure S1 plots the relationship between the aperiodic exponent and the original SPR within each diagnostic group separately. When the full samples (both AD and HC) were considered together, in each cohort separately, moderate negative relationships between original SPR and exponent were found, with r = -.488, p = .001 in Cohort 1 and r = -.621, p < .0001 in Cohort 2. This relationship was significant within the AD groups in both cohorts and within the HC group in cohort 1 (all *p'*s < .05) but did not reach statistical significance in the HC group in cohort 2 (*p* > .05).

Figure S1 further plots the relationships between the aperiodic offset and the original SPR. When the full samples were considered, moderate negative relationships were found in both cohort 1 (r = -0.380, p = .010) and cohort 2 (r = -0.457, p .002). When each diagnostic group was examined separately, AD groups in both cohorts and the HC in cohort 1 showed significant offset-ratio relationships (p's < .05), while HC from cohort 2 did not (*p* > .05) (Figure S2). Hence, taken together, these results confirm that aperiodic EEG features contribute to the previously calculated SPR.

Additionally, the relative contribution to SPR of periodic parameters describing the dominant peak within each individual spectrum was investigated. In both cohorts, correlations of peak parameters with original SPR showed that the captured oscillatory changes were primarily driven by alpha power differences and alpha center frequency, whilst bandwidth did not contribute significantly (Figure S2).

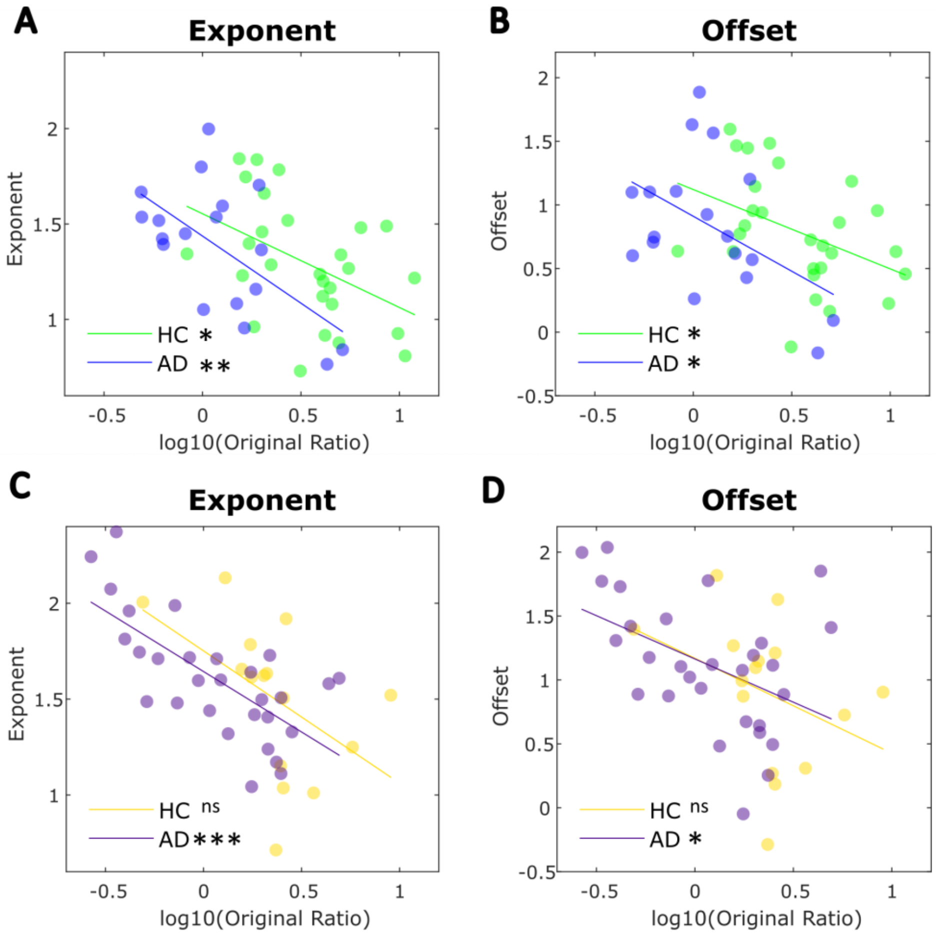


**Figure S1:** The original spectral power ratio correlates with aperiodic parameters. **A-B** Cohort 1. **C-D** Cohort 2. Pearson's correlations within each diagnostic group between the original spectral power ratio and both exponent (**A**: Cohort 1 HC r = -0.458(.016), AD r = -0.606(.008); **C:** Cohort 2 HC r = -0.491(.063), AD r = -0.697(<.0001)) and offset (**B:** Cohort 1 HC r = -0.417(.031), AD r = -0.472(.048); **D:** Cohort 2 HC r = -0.366(.179), AD r = -0.454(.013)).

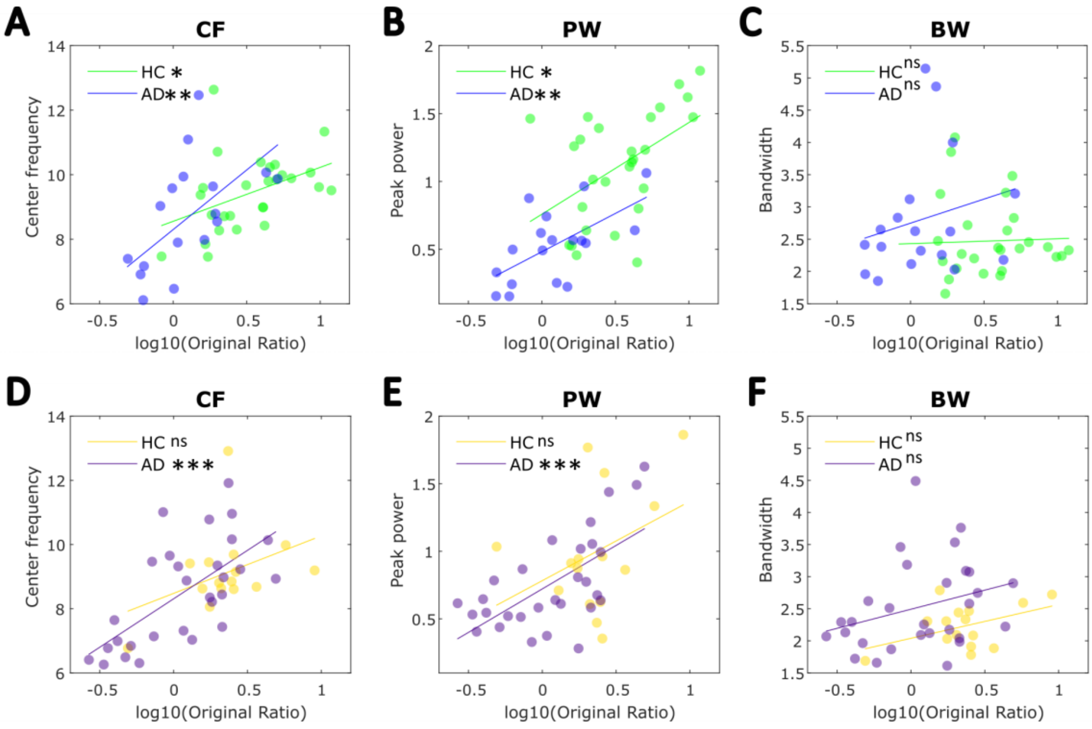


**Figure S2:** Correlations of dominant alpha (5-15 Hz) peak parameters with the original spectral power ratio. **A-C** Correlations within cohort 1 between SPR and **A** center frequency (HC r = 0.422(.028), AD r = 0.602(.008)), **B** peak power (HC r = 0.485(.010), AD r = 0.612(.007)), **C** and bandwidth (HC r = 0.035(.861), AD r = 0.229(.361)). **D-E** Cohort 2 relationships between SPR and **D** center frequency (HC r = 0.394(.146), AD r = 0.652(<.0001)), **E** peak power (HC r = 0.367(.178), AD r = 0.640 (<.0001)) , **F** and bandwidth (HC r = 0.442(.099), AD r = 0.301(.112)). When the entire sample (AD and HC) was considered together, in Cohort 1, correlations of peak parameters with the non-corrected power ratio suggested the captured oscillatory changes were primarily driven by alpha power differences (r = .702, p < .0001) followed by alpha center frequency (r = .552, p < .0001), while bandwidth did not contribute significantly (r =.031, p = .842). This pattern was similar in the second cohort, with the strongest correlations found for peak alpha power r = .580 (<.0001) and center frequency r = .607 (<.0001), while bandwidth was not associated significantly with the original SPR r = .188 (.202).

**Supplementary Section 2: Aperiodic knee model analysis**

In some subjects we noticed 'knees' (i.e., bends) in the power spectra. We therefore ran an exploratory follow up analysis using the spectral parameterization model in the knee mode. The settings for model fitting were as follows: *frequency range* 3-40 Hz (0.1 Hz resolution), *aperiodic mode* ('knee'), *peak width* ([1-12]), *maximum number of peaks* (7), *peak threshold* (1.0), and *minimum peak height* at default. The goodness-of-fit of the knee model was assessed by computing the frequency-by-frequency error for each group ($R^2$(error): Cohort 1: AD(0.997(.025)), HC(0.997(.031)); Cohort 2: AD(0.998(.027)), HC(0.997(.027)). The knee inflection point, i.e., the frequency at which the aperiodic signal begins to exponentially decrease, was calculated as $k^{\frac{1}{x}}$ (where k = knee, x = exponent). AD participants (n = 5) from the first cohort as well as one AD and one HC participant from second cohort were removed from the knee model analyses due to having implausible knee estimates (final sample sizes were as follows: Cohort 1 (AD = 13, HC = 27); Cohort 2 (AD = 28, HC = 14)).

**The 'knee' of neural power spectra differentiates between AD and HC**

Figure S3A shows the group averaged aperiodic component fitted with a knee parameter in Cohort 1. When between diagnostic group differences were considered (Figure S3B) significant differences were found in the knee frequency (F(1,37) = 15.305, p < 0.001, , $\eta^2$= 0.293), as well as offset (F(1.37) = 11.374, p = 0.002, $\eta^2$= 0.235) and exponent (F(1,37) = 11.705, p = 0.002, $\eta^2$= 0.240). As shown in Figure S3C, the knee frequency (i.e., the point at which the aperiodic fit transitions from horizontal to negatively sloped) correlated strongly with the original spectral power ratio (r = 0.807, p < 0.0001) in the 1st cohort as a whole and this relationship was statistically significant within each diagnostic group as well (HC r = 0.707(<.0001), AD r = 0.881(<.0001)).

These results were replicated in Cohort 2. Figure S3D shows the group averaged aperiodic fit with the knee included in the model for Cohort 2. The knee frequency was significantly higher in healthy controls compared to AD ($F(1,39) = 5.143$, $p = .034$, $\eta^2 = 0.189$) (Figure S3E). In line with Cohort 1, both offset ($F(1,39) = 8.718$, $p = 0.007$, $\eta^2 = 0.284$) and exponent ($F(1,39) = 10.770$, $p = 0.003$, $\eta^2 = 0.329$) showed significant between-group differences as well. As in Cohort 1, a strong positive correlation was found between the knee frequency and the original SPR ($r = 0.815$, $p < 0.0001$) which was also statistically significant when each diagnostic group was examined separately (AD: $r = 0.793$($< 0.0001$), HC: 0.750(0.002) (Figure S3F).

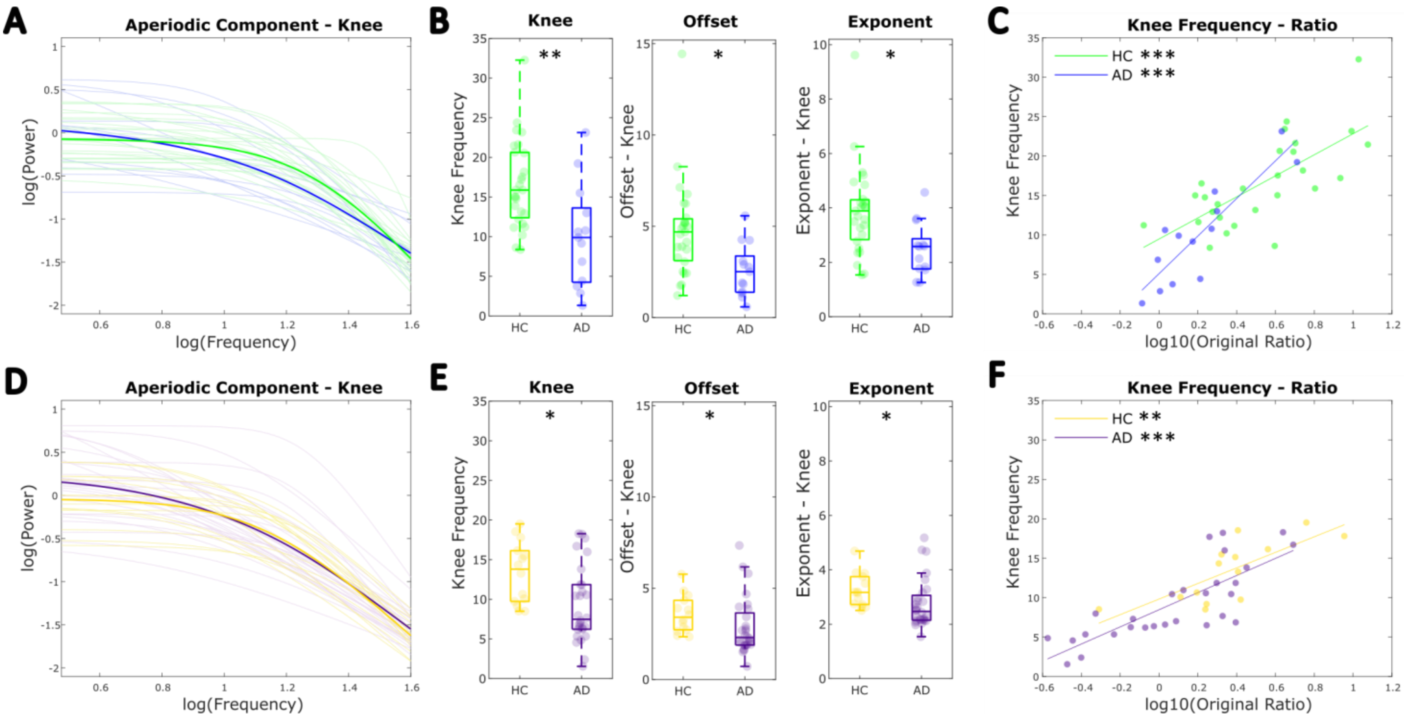


**Figure S3:** Results of analyses including an additional knee parameter in the aperiodic model. **A-C:** Cohort 1. **A:** Plots the aperiodic component of the power spectrum (bold) averaged across individuals within each diagnostic group (AD: blue, HC: green). The individual aperiodic components are also shown. **B:** Comparison of the aperiodic parameters using an ANCOVA (with *Age* included as a covariate) showed significant between-group differences in all knee frequency, offset, and exponent. **C:** Pearson's correlations between the knee frequency and the original spectral power ratio. **D-F:** Cohort 2. **D:** Group averaged aperiodic components for early AD (purple) and HC2 (yellow) groups. **E:** Between-group comparisons showed a significant difference in knee frequency, offset, and exponent. **F:** Pearson's correlations between knee frequency and SPR. *** $p < 0.0001$, ** $p < 0.001$, * $p < 0.05$, ns $p > 0.05$.

Given the significant alterations in the knee frequency in both cohorts as well as the strong relationship between knee and SPR, we also examined the between-group differences in purely oscillatory EEG measures after controlling for aperiodic signal. Figures S4A&D plot the aperiodic removed spectra for each cohort respectively. As Figure S4B shows, the difference in spectral power ratio also survived the aperiodic correction in the knee analysis: $F(1,37) = 4.217$, $p = 0.047$, $\eta^2 = 0.114$. In line with the 'fixed' analyses reported in the main text, in Cohort 1, the peak power (5-15 Hz) after removing the aperiodic activity differed significantly between the diagnostic groups, $F(1,37) = 7.923$, $p = 0.008$, $\eta^2 = 0.176$, while the corresponding centre frequency did not, $F(1,37) = 0.720$, $p = 0.402$ (Figure S4C). Additionally, the bandwidth of the identified (5-15 Hz) peak was significantly greater in AD compared to HC, $F(1,37) = 5.787$, $p = 0.021$, $\eta^2 = 0.135$ (Figure S4C). Contrastingly, no significant differences were found in Cohort 2 for the aperiodic-adjusted SPR, $F(1,39) = 0.239$, $p = 0.630$, or for peak power, $F(1,39) = 2.453$, $p = 0.132$, centre frequency, $F(1,39) = 0.354$, $p = 0.558$, and bandwidth, $F(1,39) = 3.968$, $p = 0.059$ (Figure S4E & F).

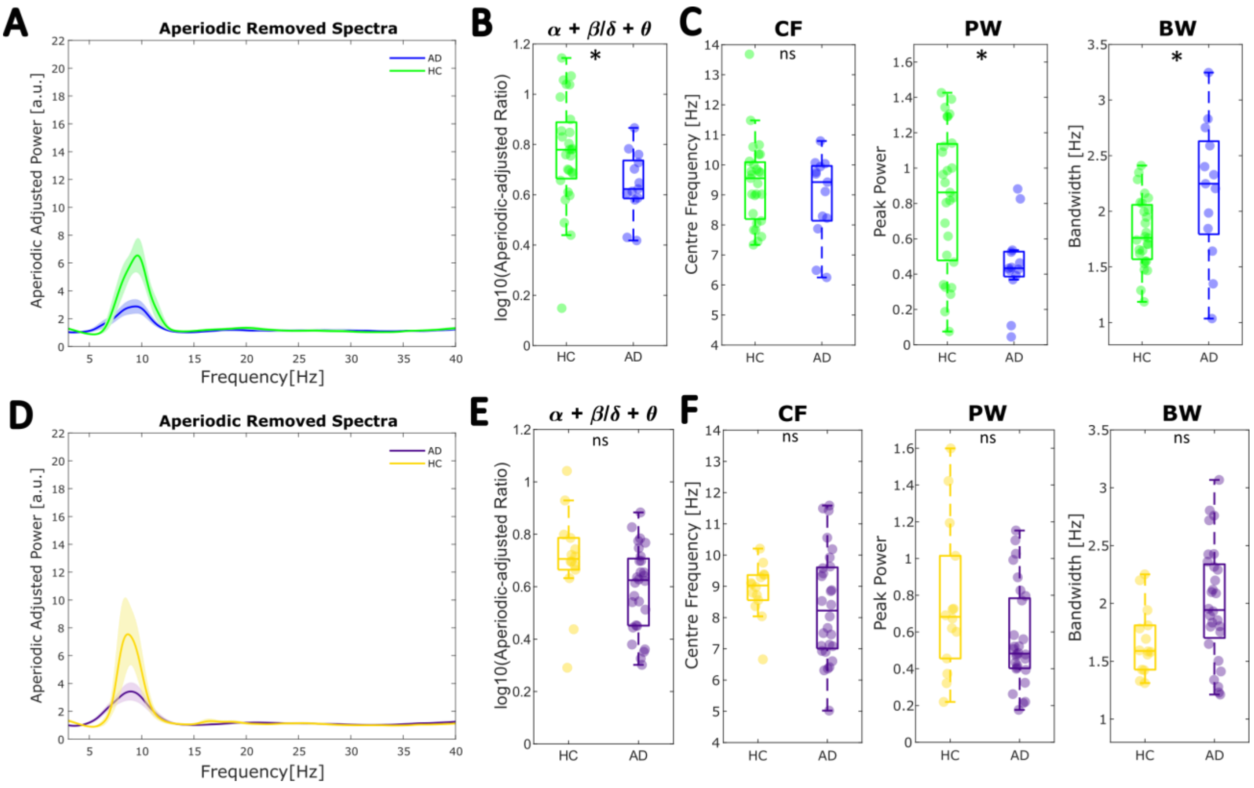

**Figure S4:** Oscillatory EEG changes in 'knee' model analysis. **A-C** Cohort 1 results. **A** The group averaged spectra after the aperiodic activity has been subtracted from the raw spectra for each participant (AD: blue, HC: green). The shaded areas represent standard error. **B** Comparison of aperiodic-adjusted SPR (log-transformed) showed a significant between-group difference. **C** Between-group comparison of periodic parameters showed power at the dominant alpha (5-15 Hz) peak is significantly reduced in AD, bandwidth is increased, whilst centre frequency does not differ. **D-F** Cohort 2 results. **D** Group averaged periodic components of the power spectrum (AD: purple, HC: yellow). Shaded area represents standard error. **E** Aperiodic-adjusted SPR computed from periodic activity did not differ significantly. **F** No significant between-group differences were found in (5-15Hz) peak parameters. * $p < .05$, ns $p > .05$.

The relationship between knee frequency and each cognitive composite measure (dementia severity, learning & memory, and executive function) were also tested. However, when age was controlled for, knee frequency did not uniquely predict any of the cognitive functions in AD or HC groups across both cohorts (all p's > .05).

While human electrophysiological recordings often knees in power spectra (especially in larger frequency ranges) (Seymour et al., 2022), currently little is known about the neurophysiological significance of the knee frequency parameter in the EEG signal. In invasive intracranial recordings, the knee frequency has been linked to neuronal timescales, which scale with cognitive functions and aging (Gao et al., 2020). However, it is unclear whether the knee represents a meaningful feature of the aperiodic signal or merely captures periodic influence on the shape of the power spectra. Our results could be consistent with the latter, as we observed very strong correlations between the knee frequency and SPR in both cohorts (1 & 2) and subtracting the aperiodic component (fitted with a knee) attenuated the between group differences in periodic features. Nevertheless, given the limited understanding of the knee frequency in non-invasive EEG recordings, we cannot interpret our results with confidence.

**Table S2**

*Results of partial correlations for the unique effects of periodic parameters on neurocognitive functions while controlling for age*

| Cognitive Function | AD Group | PW<br>*B*(*p*) | CF<br>*B*(*p*) | BW<br>*B*(*p*) |
|---|---|---|---|---|
| **Dementia Severity** | *Cohort 1* | 0.474 (.055) | 0.277 (.282) | 0.211 (.415) |
| | *Cohort 2* | 0.312 (.138) | 0.254 (.231) | 0.167 (.436) |
| **Memory & Learning** | *Cohort 1* | 0.025 (.923) | 0.396 (.115) | 0.213 (.412) |
| | *Cohort 2* | 0.161 (.453) | -0.021 (.923) | -0.271 (.199) |
| **Executive Function** | *Cohort 1* | 0.251 (.331) | 0.368 (.146) | 0.403 (.109) |
| | *Cohort 2* | **0.473 (.019)** | 0.092 (.671) | -0.122 (.570) |
| | **HC Group** | **PW** | **CF** | **BW** |
| **Dementia Severity** | *Cohort 1* | 0.230 (.259) | 0.042 (.840) | 0.116 (.572) |
| **Memory & Learning** | *Cohort 1* | 0.314 (.118) | 0.057 (.781) | <.001 (.998) |
| **Executive Function** | *Cohort 1* | -0.162 (.429) | 0.341 (.088) | 0.238 (.243) |

*Note:* We were unable to analyse relationships to neuropsychological function in HC group in cohort 2 due to low numbers of participants who underwent cognitive testing.

**Table S3**

*Results of partial correlations for the unique effects of periodic and aperiodic measures on neurocognitive functions while controlling for age within the HC groups*

| Cognitive Function | HC Group | Original Ratio<br>*B*(*p*) | Aperiodic-adjusted Ratio<br>*B*(*p*) | Exponent<br>*B*(*p*) | Offset<br>*B(p)* |
|---|---|---|---|---|---|
| **Dementia Severity** | *Cohort 1* | 0.266 (.189) | 0.203 (.320) | 0.044 (.831) | -0.147 (.475) |
| **Memory & Learning** | *Cohort 1* | 0.237 (.244) | -0.012 (.952) | -0.325 (.106) | -0.287 (.155) |
| **Executive Function** | *Cohort 1* | 0.190 (.352) | 0.220 (.281) | -0.272 (.179) | -0.316 (.116) |

*Note:* We were unable to analyse relationships to neuropsychological function in HC group in cohort 2 due to low numbers of participants who underwent cognitive testing.
